# Excitatory and inhibitory neurons in the dorsal periaqueductal gray encode decisions to assess and escape natural threats

**DOI:** 10.64898/2026.02.16.706076

**Authors:** Irene P. Ayuso-Jimeno, Sofia Torchia, Valeria Mussetto, Taddeo Salemi, Cornelius T. Gross

**Affiliations:** Epigenetics & Neurobiology Unit, European Molecular Biology Laboratory (EMBL), EMBL Rome, Via Ramarini 32, 00015 Monterotondo (RM), Italy; Joint PhD degree program, European Molecular Biology Laboratory and Faculty of Biosciences, Heidelberg University, 69117 Heidelberg, Germany

## Abstract

Prey species are able to engage hardwired neural pathways to rapidly escape from an imminent predator attack. However, when predator threat is less probable they typically show a stereotypical sequence of approach toward the threat aimed at gathering more information, followed by escape to safety when the threat threshold is reached. The brainstem dorsal periaqueductal gray (dPAG) is required for the expression of escape behavior to predator threats and stimulation of dPAG elicits goal-directed flight. However, *in vivo* neural recordings in dPAG have identified separate populations of neurons that are tuned to either the approach or escape phase of the behavior suggesting that the structure may also be involved in threat assessment. The genetic identity and connectivity of these *Assessment*+ and *Escape*+ neurons have not been defined, although optogenetic activation of glutamatergic, but not GABAergic neurons elicits high-speed flight, suggesting that *Escape*+ neurons might be exclusively excitatory in nature. Moreover, it is not clear whether non-predator threats such as those elicited by conspecific or other animate threats are encoded by independent or overlapping neurons in dPAG. Here we report the activity pattern of ensembles of glutamatergic and GABAergic dPAG neurons during approach and escape from predator, social, and prey threats. Unexpectedly, we found that both glutamatergic and GABAergic neurons harbor *Assessment*+ and *Escape*+ neurons, suggesting that both cell-types are engaged in the approach-to-avoidance transition. Consistent with the functional involvement of both cell-types in approach-to-avoidance behavior, optogenetic activation of GABAergic cells elicited a reduction of risk assessment behavior towards the predator. Finally, we found that exposure to predator, social or prey threat recruited largely overlapping neurons in dPAG, demonstrating a convergence of threat processing in this structure. These findings point to a tightly coordinated role for dPAG excitatory and inhibitory neurons in the generalized control of innate threat assessment and avoidance behavior.

## Introduction

The periqueductal gray (PAG) is a brain nucleus located in the rostral brainstem that has been linked to the control of defensive and other innate behaviors across animal species. Lesion and electrical stimulation studies in laboratory animals have allowed for the anatomical and functional segmentation of PAG into a dorsal and ventral division with the former associated with the production of freezing and flight responses to threat, defensive and appetitive vocalization, and activation of the sympathetic autonomic system, while the latter has been associated with the promotion of immobility and the suppression of ongoing appetitive behaviors, analgesia, and activation of the parasympathetic autonomic system (Amano et al., 1982; Bittencourt et al., 2004; Johansen et al., 2010; Gross & Canteras, 2012). The dorsal PAG is further divided into dorsomedial, dorsolateral, and lateral subdivisions which, although they have distinct efferent and afferent projections, have not yet been assigned distinct functional roles in behavior or autonomic regulation (Beitz et al., 1982; Carrive, 1993; Vianna and Brandão, 2003; Fillinger et al., 2017). Importantly, electrical stimulation of dorsal PAG in humans elicits aversive feelings and the sensation of being chased, suggesting a conserved role in the promotion of defensive states also in humans (Nashold et al. 1969; Amano et al. 1982).

A functional cellular and circuit architecture of dorsal PAG is emerging in which stimulation of glutamatergic neurons in dorsal PAG promotes freezing at low intensity and flight at high intensity (Tovote et al. 2016, Deng et al. 2016, Evans et al. 2018, Tsang et al. 2023). The basis for the transition in evoked responses from freezing to flight remains unknown, but suggests a non-linearity in circuit activity either at the level of dPAG or its downstream brainstem targets (Tsang et al. 2023). Stimulation of GABAergic neurons, on the other hand, does not elicit defensive behavior (Tsang et al., 2023) and recent evidence shows that they can inhibit looming stimulus-evoked flight behavior (Stempel et al., 2024) suggesting they might act as part of a tonic inhibitory circuit that receives primarily local inputs (Franklin et al., 2017; but see Wu et al. 2024). Such a push-pull circuitry controlling defensive behavior is compatible with the identification of two distinct classes of neurons in single unit recording studies in mice approaching and fleeing from a predator. *Assessment*+ neurons gradually increased their firing as the animal approached the threat, but abruptly stopped firing at the onset of flight, while *Flight*+ neurons were suppressed during approach and showed a sudden increase in firing at flight onset (Deng et al. 2016; Masferrer et al. 2020). These data suggest a model in which *Flight*+ cells are excitatory neurons that promote escape, while *Assessment*+ cells are inhibitory neurons that suppress locomotion during approach to a threat, and permit flight by shutting off at escape onset. However, an alternative model argues that local feedback inhibition has a role in the abrupt switch in activity between *Assessment*+ and *Flight*+ cells at escape onset (Rahy et al. 2022). If true, this model predicts that local inhibitory neurons should have similar *Assessment*+ and *Flight*+ populations as excitatory neurons.

A second question arises concerning the convergence of threat processing in dPAG. Dorsal PAG receives its primary defense-related inputs from the medial hypothalamic defensive network (Beitz 1982, Marchand & Hagino 1983, Canteras 2002; Gross & Canteras 2012; Silva et al. 2013) where the processing of predator and social threats is anatomically segregated (Motta et al. 2009; Silva et al. 2013). While dPAG has been shown to be required for the production of defensive behaviors to both predator and social threats (Silva et al. 2013), it is not known if this occurs via the same or different neurons in dPAG. Here we set out to test these hypotheses. We found that excitatory and inhibitory neurons show similar encoding of approach-avoidance behavior with both showing prominent *Assessment*+ and *Escape*+ (formerly referred to as *Flight*+) response profiles. Serial exposure to predator, social, and prey threats activated largely overlapping *Assessment*+ and *Escape*+ cells. These findings argue for the convergent control of approach-avoidance behavior in dPAG and suggest that local feedback inhibition may play a role in the decision to escape threat.

## Results

### Both excitatory and inhibitory dPAG neurons encode assessment and escape

To investigate the genetic identity of *Assessment*+ and *Escape*+ neurons, we recorded calcium transients via GCaMP6f fluorescence imaging through a miniaturized microscope while mice approached and escaped from a rat, a natural murine predator. Our experimental mice were exposed to a laboratory rat in a custom-made behavioral apparatus consisting of a chamber, a corridor and a second chamber in which the rat was placed. A plexiglass separator with a small opening in the bottom was placed between the corridor and the rat chamber to prevent the rat from attacking the mouse, while allowing for sniffing through the opening (**Figure 1A**, top). In this setting, the mouse repeatedly approached the rat at low velocity, assessing the threat and subsequently escaped at high velocity moving away from the rat (**Figure 1A**, bottom; **Figure 1B**; **Figure S1**). Miniscope recordings were performed in *Vglut2*::Cre and *Vgat*::Cre mice in order to monitor activity in the principal dPAG inhibitory and excitatory cell-types (Vaughn et al., 2022). dPAG was reached with a chronically implanted GRIN lens (**Figure 1C**, left, **Figure S1**) which allowed for the recording of single-unit GCaMP6f-derived fluorescence (**Figure 1C**, right). This experimental setting allowed for the identification of putative single neuronal somata in videos acquired from both *Vglut2*::Cre and *Vgat*::Cre mice after preprocessing of the miniscope videos (see **Materials & Methods** section; **Figure 1D**; **Figure S1**). Next, the average fluorescence of all pixels belonging to each unit over time was extracted and time-locked to the onset of escape behavior, allowing for the examination of calcium transients surrounding risk assessment and escape behavior as well as behavior metrics like velocity (**Figure 1E**). We found *Vglut2^+^* neurons whose activity consistently rose during risk assessment and dropped at escape onset across trials (*Assessment*+, **Figure 1F**). Likewise, other neurons showed an opposite pattern of activity, with their calcium signal dipping before escape and rising at escape onset (*Escape*+, **Figure 1G**). We assume that these neurons correspond to the previously reported *Assessment*+ and *Flight*+ neurons, respectively, measured by *in vivo* single-unit electrophysiology (Deng et al. 2016; Masferrer et al., 2020). Next, we examined the activity of *Vgat*^+^ neurons in relation to assessment and escape behavior (**Figure 1H**). Surprisingly, *Vgat*^+^ neurons also showed populations with consistent *Assessment*+ and *Escape*+ response properties across assessment and escape trials (**Figure 1I-J**).

**Figure 1.**
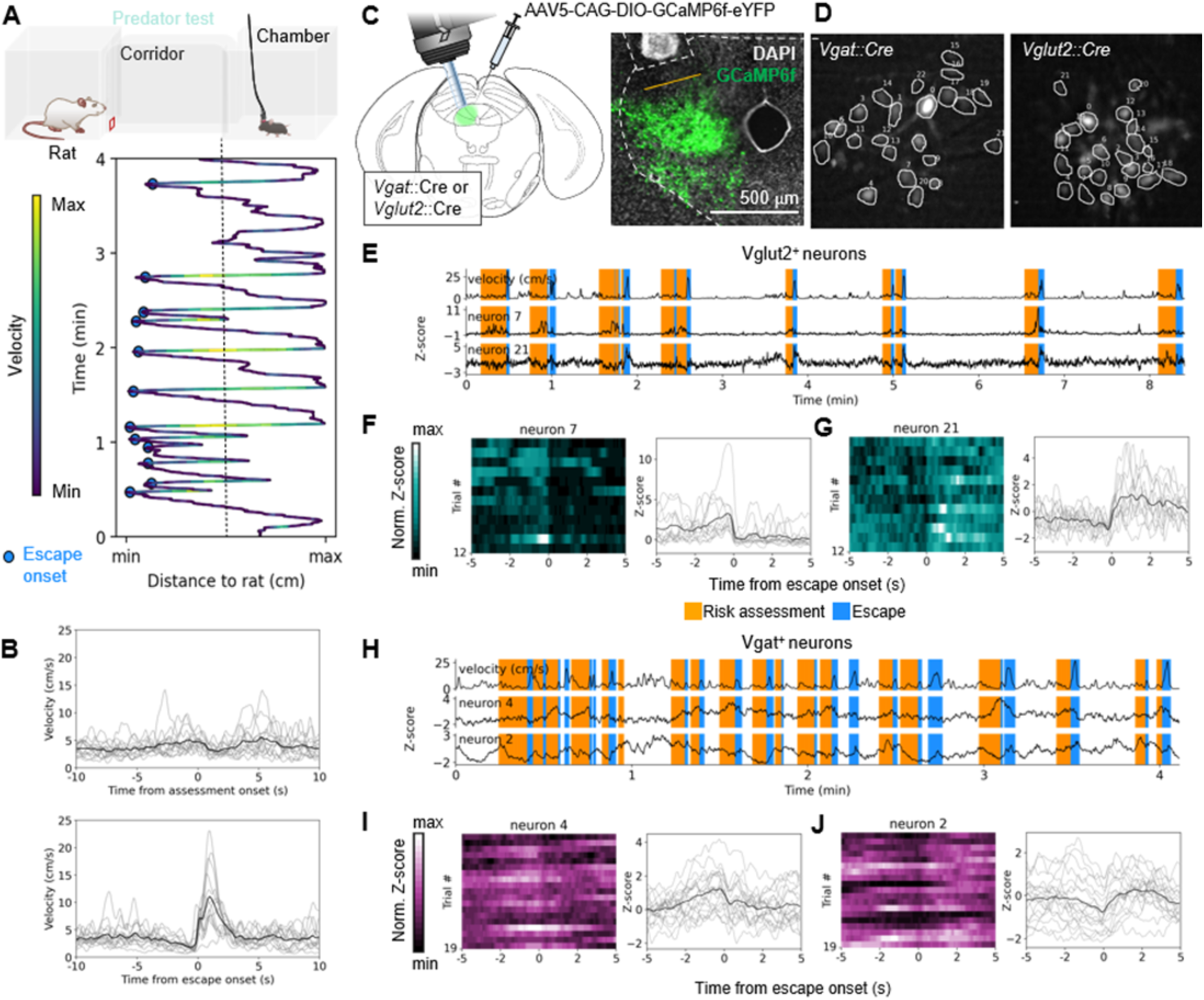
Calcium imaging of *Vglut2*+ and *Vgat*+ dPAG neuron activity during predator assessment and escape behavior. (**A**) Top: Sketch representing the behavioral apparatus composed of two chambers and a corridor. A rat is placed in a chamber separated from the corridor by a wall with a small opening (red) which allows for interaction but impedes aggression. Bottom: distance to the rat over time for one exemplary mouse; color code indicates velocity. Slow velocity assessment followed by high velocity escape is shown repeatedly. (**B**) Velocity of the mouse centered around assessment onset (top) and escape onset (bottom). (**C**) Left: Schematic of virus infection location and GRIN lens placement in dPAG. Right: histology section showing exemplary GRIN lens tract and eYFP fluorescence (green) which is a proxy for GCaMP6f expression. (**D**) Maximum projections of miniscope fields of view (FOV) with regions of interest overlayed for an exemplary *Vgat*::Cre (left) and *Vglut2*::Cre (right) mouse line. (**E**) Top: velocity of the center of mass over time for a *Vglut2*::Cre mouse. The overlaid color indicates risk assessment (orange) or escape (blue) behavior. Middle: Example neuron (neuron 7) belonging to the same *Vglut2*::Cre mouse showing activity rising during risk assessment behavior. Bottom: example neuron (neuron 21) showing activity increasing during escape behavior. (**F**) Left: heatmap of all assessment and escape trials for neuron 7 centered around escape onset. Right: average activity over trials showing *Assessment*+ properties (**G**) Left: heatmap of all trials for neuron 21 centered around escape onset. Right: average activity over trials showing *Escape*+ properties. (**H-J**) as E-G for a *Vgat*::Cre mouse. Significant correlations were calculated by comparing the average slope for a given neuron at escape onset with 1000 randomly shuffled average slopes along the entire trace of the neuron. Data are presented as mean (black) and individual trials (grey).

To further explore the engagement of dPAG excitatory and inhibitory neurons during risk assessment and escape behavior we examined the activity of the *Vglut2^+^* population (111 neurons, 6 mice) in more detail at risk assessment onset (**Figure 2A**, left). We observed that 16% of *Vglut2^+^* neurons showed activity that dropped during risk assessment behavior (**Figure 2B** and **C**, top), and 21% showed activity that rose during this period (**Figure 2B**, bottom; **Figure 2D**). Interestingly, these two classes showed opposite activity patterns at escape onset (**Figure 2A**, right, **B** and **C** bottom) showing that *Assessment*^+^ neurons are active during risk assessment and deactivated at escape, and *Escape*^+^ neurons are silent during risk assessment and activated during escape. Escape behavior elicited a larger number of correlated neurons for both classes, with 40% *Assessment*^+^ and 33% *Escape*^+^ neurons (**Figure 2D**). Next, we examined the activity of the *Vgat^+^* population (166 neurons, 8 mice). To our surprise, *Vgat^+^* neurons showed similar activity patterns to *Vglut2^+^* neurons around risk assessment and escape behavior (**Figure 2E-G**). Moreover, *Escape*^+^ neurons were more abundant in GABAergic than glutamatergic neurons, with 10% *Assessment^+^* and 25% *Escape^+^* neurons detected when locked with assessment onset, and 17% and 44% detected, respectively, when locked with escape onset (**Figure 2H**). However, we noticed that their temporal dynamics differed, with GABAergic neurons showing slower responses.

**Figure 2.**
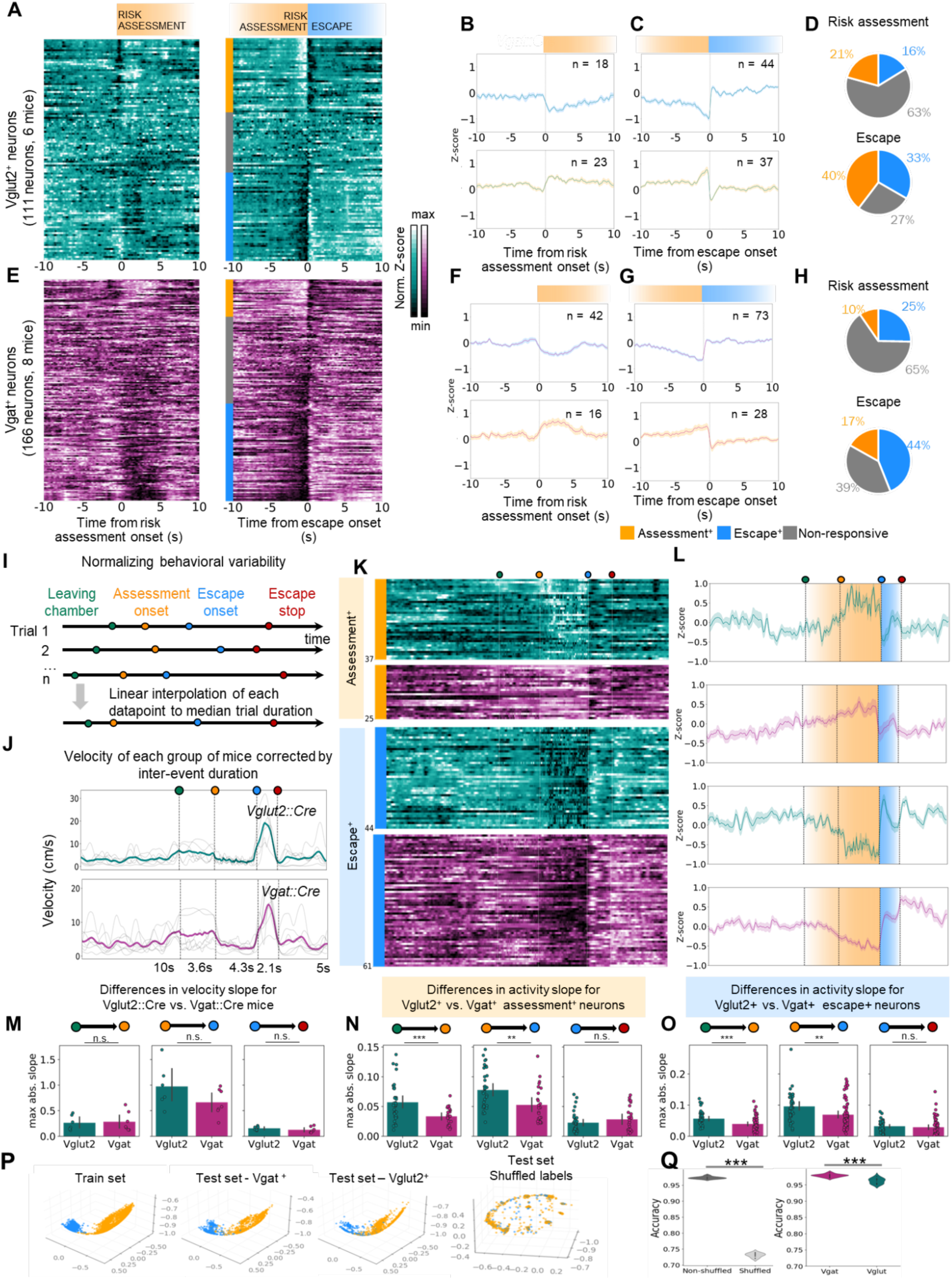
Matching dynamics of neuronal activity and behavior in excitatory and inhibitory neurons. (**A**) Heatmaps showing normalized glutamatergic activity recorded in 111 neurons across 6 mice locked to risk assessment onset (left) and escape onset (right). Neurons are sorted according to responsiveness to escape onset (blue: *Assessment*^+^ neurons; orange: *Escape*^+^ neurons; grey: non-resonsive neurons). (**B**) Average activity of glutamatergic neurons is categorized as *Assessment^−^* (top) and *Assessment*^+^ (bottom). (**C**) Average activity of *Escape*^+^ (top) and *Assessment*^+^ neurons (bottom) locked to escape onset. (**D**) Proportions of categorized glutamatergic neurons around risk assessment (top) and escape (bottom) onset. (**E-H**) As A-D, but for GABAergic neurons (N = 166 neurons, N = 8 mice). (**I**) Schematic illustrating the rationale behind linear time warping (LTW) of behavioral and activity time series. (**J**) Maximum absolute slope of velocity of the center of mass for *Vglut2*::Cre and *Vgat*::Cre mice in the behavioral windows indicated in I. (**K**) Heatmaps of activity of *Assessment*^+^ and *Escape*^+^ glutamatergic (teal) and GABAergic (violet) neurons during risk assessment and escape behavior after LTW. (**L**) Average activity of neurons displayed in M. (**K**) top: maximum absolute slope of Z-scored activity for *Assessment*^+^ *Vglut2*^+^ and *Vgat*^+^ neurons in warped behavioral windows. Bottom: maximum absolute slope of Z-scored activity for *Escape*^+^ *Vglut2*^+^ and *Vgat*^+^ neurons in warped behavioral windows. Significant correlations were calculated by comparing the average slope for a given neuron at escape onset with 1000 randomly shuffled slopes along the entire trace of the neuron. Statistical difference in slope values (**M-P**) was calculated with a paired t-test (***p < 0.001, **p < 0.01, *p < 0.05). Data are presented as mean +/- SEM. (**P**) From left to right: Embedding obtained by training multi-session Cebra Behavior on 70% of each mouse’s neural activity and behavioral labels acquired during the predator experimental test; embedding obtained by testing the previous model on unseen data acquired from *Vgat*+ mice during the same experimental test; embedding similarly obtained by testing the previous model on *Vglut2*+ data from the same experimental test; multi-session CEBRA Behavior embedding obtained by training the model on 70% of each mouse’s neural activity and behavioral labels acquired during the predator experimental test after randomly shuffling the behavioral labels. (**Q**) Left: comparison between prediction accuracy of behavioral labels using KNN of 1) 10 different multi-session CEBRA Behavior models trained and tested on 10 separate random train-test splits (all mice, predator test) and 2) 10 different multi-session CEBRA Behavior models trained and tested with shuffled behavioral labels (same mice, predator test, new random train-test split). Right: prediction accuracy of behavioral labels using KNN of 10 different multi-session CEBRA Behavior models trained on 10 separate randomly extracted training data (all mice, predator test) and tested only on *Vgat*+ mice test data or on *Vglut2*+ mice test data. Statistical significance was tested with Wilcoxon rank-sum (**p < 0.001, **p < 0.01, *p < 0.05).

To better visualize correlations between neural activity and the onset and offset of defensive behaviors we normalized the duration of peri-event windows using linear time warping (Stempel et al., 2024). We selected the onset of risk assessment, the entry into the area of the corridor near the rat, the onset of escape, and the end of escape for each trial, and interpolated the time series to the median length of the trial-averaged Z-scored values of all windows for all mice (**Figure 2I**, see **Materials & Methods**). First, we time-warped the velocity series for *Vgat*::Cre and *Vglut2*::Cre mice separately (**Figure 2J**) finding that traces did not differ significantly in the velocity slopes for any of the studied windows (**Figure 2K**). This data suggested that the overall behavior of the two Cre lines did not differ significantly, as expected (see **Figure S1** for non-time-warped data). We next proceeded to time warp the neural data of the previously classified *Assessment^+^* and *Escape^+^* responsive neurons from each population (**Figure 2L-M**). As suggested by the non-interpolated data, the slopes of *Vgat^+^* and *Vglut2^+^* neurons differed significantly for the first two analyzed time windows, leaving the chamber-to-risk assessment onset and risk assessment onset-to-escape onset. Both *Assessment^+^* and *Escape^+^* neuron activity in the *Vgat*^+^ population ramped up or decreased with a smaller maximum absolute slope than for those in the *Vglut2*^+^ population, indicating slower dynamics for the risk assessment phase of the behavior (**Figure 2N**).

Next we investigated the relationship between the activity of the entire population of neurons and behavior, by applying CEBRA (Schneider et al., 2023) to the *Vglut2*+ and *Vgat*+ population, focusing exclusively on time points corresponding to either risk assessment or escape behavior. We trained CEBRA models using the calcium traces from both *Vglut2*+ and *Vgat*+ neurons, and evaluated behavior classification accuracy across trials (**Figure 2P, S3C**). When tested across all mice, the model reliably separated risk assessment from escape epochs, indicating a robust encoding of behavioral state in the population activity. Importantly, this performance held when we trained and tested on *Vglut2*+ and *Vgat*+ mice separately, suggesting that both excitatory and inhibitory populations contain sufficient information to distinguish approach-avoidance behavioral states. To validate these findings, we benchmarked the classification results against a control condition in which behavioral labels were randomly shuffled. Here, prediction accuracy dropped dramatically, confirming that the high performance observed in the original models was not due to chance. To further quantify this effect, we cross-validated these results by iterating the training and testing procedure ten times for both the original and shuffled datasets. The difference between the shuffled and non-shuffled conditions (**Figure 2Q, left**) was statistically significant, reinforcing the conclusion that neural activity patterns in both *Vglut2*+ and *Vgat*+ populations reliably encode behavioral states. Additionally, we cross-validated the classification accuracy of CEBRA separately on *Vglut2*+ and *Vgat*+ populations (**Figure 2Q, right**). While both populations yielded high accuracy, the difference between them was statistically significant, suggesting subtle differences in how excitatory and inhibitory neurons represent behavioral state (**Figure 2Q**). To test whether this difference in accuracy was dependent on the training set, we also trained CEBRA models exclusively on *Vglut2*+ or *Vgat*+ neurons and tested them on the same population with the same cross-validation technique (**Fig. S3A,B,I**). These models reproduced the original accuracy differences, indicating that the observed disparity is intrinsic to the population-specific encoding rather than a result of cross-population training.

### Inhibitory neurons modulate defensive and exploratory behavior

Previous studies have shown that optogenetic activation of *Vglut2*^+^ dPAG neurons elicited flight behavior (Deng et al., 2016; Evans et al., 2018; Tsang et al., 2023). Optogenetic activation of *Vgat*+ dPAG neurons, however, did not elicit flight (Tsang et al. 2023). To test whether inhibitory neurons in dPAG might nevertheless block defensive behaviors elicited by a predator, we optogenetically activated *Vgat*^+^ dPAG neurons in the presence of a rat. We infused adeno-associated virus (AAV) carrying a Cre-dependent ChR2 or eYFP (control animals) bilaterally into the dPAG of *Vgat*::Cre mice and optic fibers were implanted above the AAV infection sites (**Figure 3A**, **Figure S2A**). Blue light (465 nm) was delivered to the animals for alternating one minute intervals for a total of six minutes. Light stimulation did not change overall locomotor behavior (**Figure 3B**) measured as velocity, time spent in the corridor, or number of entries in the risk area (**Figure 3C**) confirming earlier findings that inhibitory neuron activation does not induce flight (Tsang et al. 2023). When looking closely at risk assessment and escape events, however, optogenetic stimulation of *Vgat*^+^ dPAG neurons reduced animal velocity during both risk assessment and escape (**Figure 3D-E**). These data suggest that the quality of risk assessment and escape behavior is modulated by inhibitory neuron activity. To test this in a more unbiased manner we automatically categorized behavior using keypoint-MoSeq (Weinreb et al., 2024), an unsupervised machine learning method for animal behavior annotation that analyzes DeepLabCut extracted poses and segments them into syllables. In our datasets, 95% of the examined frames were represented by the first 16 syllables (**Figure S2B**). To decode the behavior of the most frequently observed syllables we manually annotated key behaviors in our videos focusing on risk assessment, retraction, escape, rearing and jumping (see **Materials & Methods** for definitions of each behavior). Notably, two syllables, 3 and 10, appeared to reflect rearing behavior, syllable 12 represented escape behavior, and syllable 13 represented risk assessment behavior (**Figure 3F-G**). After unsupervised categorization of behavior we proceeded to statistically test the differences in time spent in a given syllable with and without optogenetic stimulation of *Vgat*^+^ dPAG neurons. Amongst significant syllables (3,4,7,10,11,13,15; **Figure S2C**) *Vgat*^+^ dPAG stimulation induced a significant increase in the time spent in rearing behavior and a significant decrease in the time spent in risk assessment behavior when compared to control animals (**Figure 3H**). This finding was confirmed by a *post hoc* analysis of the manually annotated time spent in rearing and risk assessment behavior (**Figure 3I**). Interestingly, although stimulation of Vgat⁺ dPAG neurons significantly reduced the peak speed of escape, it did not change the overall time spent escaping (**Figure S2D**). This suggests that activating the inhibitory dPAG network decreases defensive, risk-assessment behaviors toward a predator and promotes more exploratory behaviors, such as rearing.

**Figure 3.**
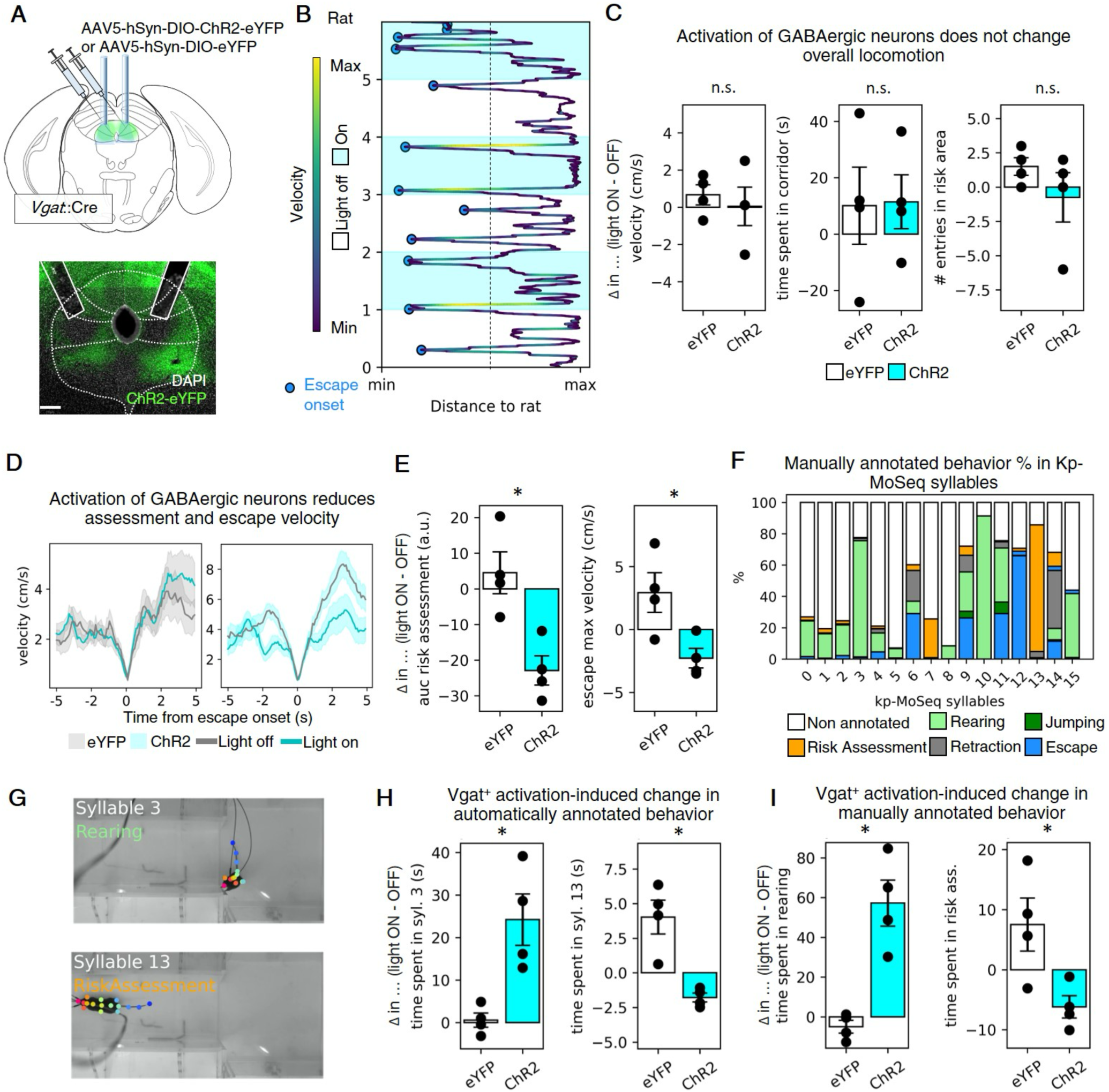
Inhibitory neurons modulate defensive and exploratory behavior. (**A**) Top: Schematic of virus infection location and optic fiber placement in dPAG. Bottom: histology section showing exemplary optic fibre tracts and for ChR2-eYFP presence. (**B**) Distance to the rat over time color-coded by velocity for a representative ChR2 expressing animal (white, light on; blue, light off). (**C**) Effect of ChR2 stimulation of *Vgat2*^+^ PAG cells on locomotion measured as the light-induced difference in velocity (light on - light off) for ChR2 vs eYPF (control) expressing mice (left), in time spent in the corridor (middle), and number of entries in risk area, defined as the half of the corridor closer to the rat (right). (**D**) Velocity time-locked to escape onset for eYFP (left) and ChR2 (right) animals. (**E**) Light-induced difference in area under the curve (AUC) for a risk assessment window (left) and peak escape velocity (right) for eYFP and ChR2 mice. (**F**) Composition of kp-moSeq syllables compared to manually annotated behavior. (**G**) Representative video frames corresponding to syllable 3/rearing (top) and syllable 13/risk assessment (bottom). (**H**) Light-induced difference in time spent in kp-MoSeq syllable 3 (left) and 13 (right) for eYPF and ChR2 mice. (**I**) Light-induced difference in time spent in manually annotated rearing (left) and risk assessment (right). Statistical significance was tested in panels C, E, H and I with Wilcoxon rank-sum, eYFP: N = 4, ChR2: N = 4. ***p < 0.001, **p < 0.01, *p < 0.05. Data are presented as mean +/-SEM.

### Convergent encoding of predator, social, and prey threat in dPAG

Although dPAG has been shown to be required for defensive responses to both predator and social threats (Silva et al. 2013) it is not known if these are mediated by the same cells within this brain region. To test this, prior to the above described rat exposure test, we sequentially exposed mice to an aggressive conspecific (CD1 ex-breeder male mouse) and a live cockroach – a novel prey known to elicit defensive responses (Rossier et al. 2021), and used miniscope calcium imaging to compare behavior-locked neuronal responses (**Figure 4A,B**). Average mouse velocity at escape onset followed a similar dynamic across the three tests, with a reduction of velocity shortly before escape onset (risk assessment) followed by a rapid increase in velocity (escape) followed by a decrease returning to baseline velocity (escape termination). However, the magnitude of escape velocity differed, with predator-induced escape trials being the ones reaching the highest peak velocity, followed by cockroach-induced and social induced escape. Moreover, prey-induced escapes were terminated earlier than social and rat-induced escapes (**Figure 4C**). As in the predator test (**See Figure 1A**), mice repeatedly approached and escaped an aggressive conspecific, with slow velocity locomotion towards the CD1 mouse followed by a change in direction and high velocity escape. Similarly, mice also repeatedly approached and escaped the cockroach (**Figure 4D**). Next, we proceeded to look at escape-locked neural activity in the *Vglut2^+^*population (111 neurons, 6 mice). We found that, similarly to what we observed in the rat test, a class of dPAG excitatory neurons showed opposing patterns of activity locked to the onset of escape triggered by both the aggressive conspecific and the prey. One third (34%) of *Vglut2^+^* neurons showed activity that decreased during risk assessment behavior and rose during escape and 19% showed activity that rose during risk assessment and dropped during escape triggered by an aggressive conspecific (**Figure 4E,G, top**). Instead, in the prey exposure test, 24% of *Vglut2^+^* neurons showed *Escape^+^* activity vs. 14% showed *Assessment^+^* activity (**Figure 4F,H**, top). We next examined the activity of the *Vgat^+^* population (166 neurons, 8 mice) upon exposure to the three threats. *Vgat^+^* neurons showed similar activity patterns to *Vglut2^+^* neurons around risk assessment and escape behaviors in both the aggressive conspecific and prey exposure tests (Social test: 31% *Escape^+^* and 13% *Assessment^+^* neurons; Prey test: 37% *Escape^+^* and 14% *Assessment^+^* neurons. **Figure 4E-H**, bottom). These data suggest that, as previously reported (Silva et al. 2013, Reis et al. 2021a, Reis et al. 2021b), dPAG neurons are involved in risk assessment and escape behavior across biological threats.

**Figure 4.**
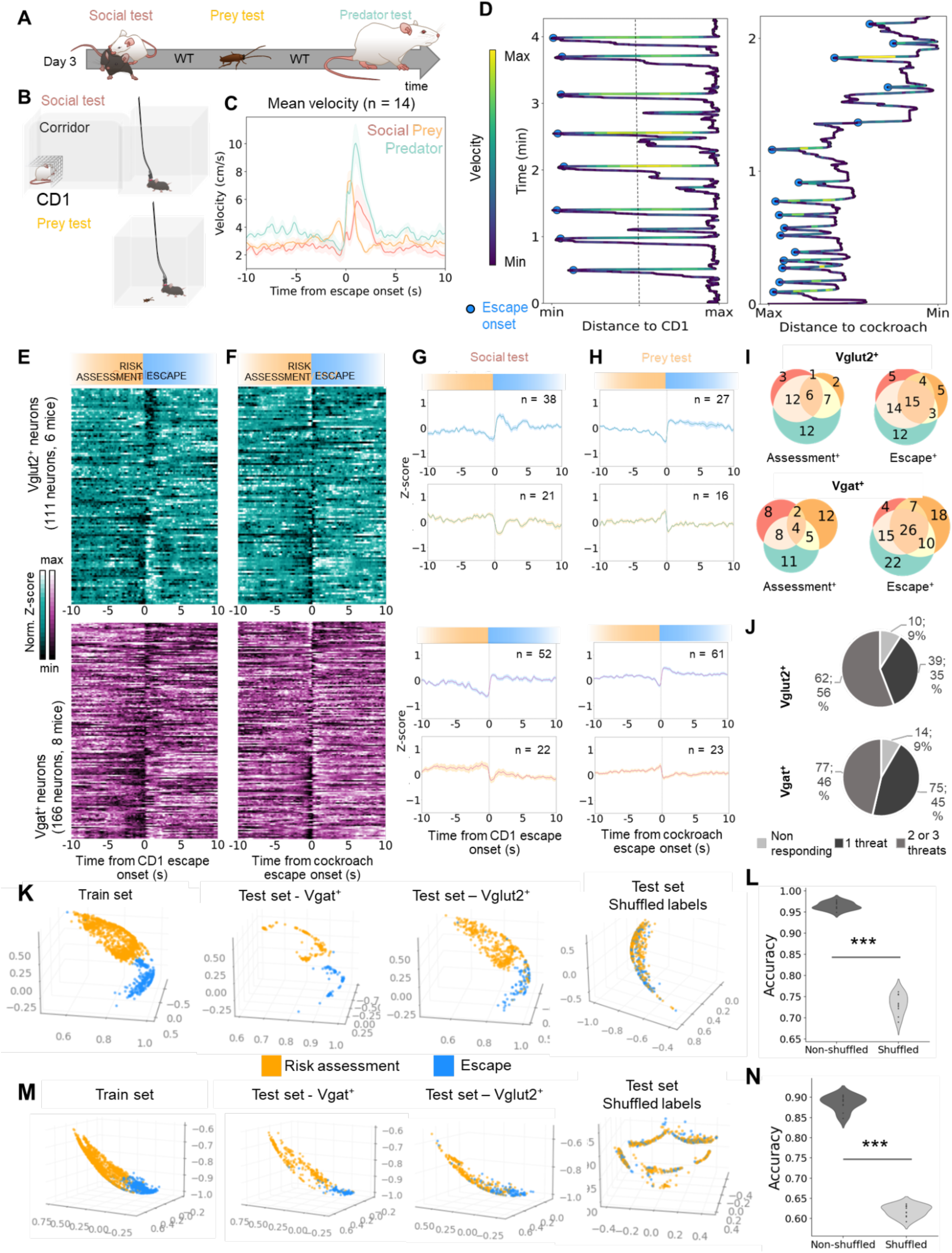
Convergent encoding of predator, social, and prey threat in excitatory and inhibitory dPAG neuron assemblies. (**A**) Sketch representing the behavioral task design, where mice were exposed sequentially to an aggressive conspecific (CD1 mouse), a prey (cockroach) and a predator (rat). At least 5 minutes separated tests to allow GCaMP photobleaching to recover. Mice remained attached to the miniscope to facilitate accurate tracking of neurons across tests. (**B**) Sketches representing the behavioral arenas in the social test (top) and prey test (bottom). (**C**) Velocity of the mouse centered around escape onset for the three behavioral tests. (**D**) Distance to the CD1 (left) or cockroach (right) over time for a representative mouse; color code indicates velocity. Slow-velocity assessment followed by high-velocity escape was shown repeatedly. (**E**) Top: heatmaps showing normalized glutamatergic activity recorded in 111 neurons across 6 mice locked to risk escape onset for the social test. Neurons are sorted according to responsiveness to escape onset in the rat test (see Fig. 2) to allow for visual comparisons of activity across tests. Bottom: as top, for GABAergic neurons (166 neurons). (**F**) as E, for the prey test. (**G**) Top: average activity of glutamagergic neurons categorized as *Assessment^+^* (orange) and *Escape^+^* (blue) in the social test. Bottom: as top, for GABAergic neurons. Data are presented as mean +/- SEM. (**H**) as G, for the prey test. (**I**) Venn diagrams representing activity overlap across tests (social: red; prey: yellow; predator: green) for glutamatergic neurons (top) and GABAergic neurons (bottom). (**J**) Pie charts representing the summary of responsiveness and activity overlaps for excitatory (top) and inhibitory neurons (bottom). (**K**) From left to right: Embedding obtained by training multi-session CEBRA Behavior on 70% of each mouse’s neural activity and behavioral labels acquired during the social test; embedding obtained by testing the previous model on unseen data acquired from *Vgat*+ mice during the same experimental test; embedding similarly obtained by testing the previous model on *Vglut2*+ data from the same experimental test; multi-session CEBRA Behavior embedding obtained by training the model on 70% of each mouse’s neural activity and behavioral labels acquired during the social test after randomly shuffling the behavioral labels. (**L**) Comparison between prediction accuracy of behavioral labels using KNN of 1) 10 different multi-session CEBRA Behavior models trained and tested on 10 separate random train-test splits (all mice, for social test and 2) 10 different multi-session CEBRA Behavior models trained and tested with shuffled behavioral labels (same mice and experimental tests, new random train-test splits). (**M,N**) as K,L for prey test. Statistical significance was tested with Wilcoxon rank-sum (*p < 0.001, p < 0.01, *p < 0.05).

To assess whether the same or different assemblies of excitatory and inhibitory neurons responded to each threat, we analyzed the overlap in tuning across tests. More than half (56%; 62/101) of the responsive *Vglut2^+^* neurons (111 in total) responded to the onset of escape in a shared manner across two or three threats and 35% responded to a single threat (39/111), whereas 46% (77/166) of the responsive *Vgat^+^*neurons respond to the onset of escape in a shared manner, and 45% responded to one threat (75/166) (**Figure 4I,J**).

To further validate our findings, we used CEBRA-based analysis on data from the aggressive conspecific and prey exposure tests. Consistent with our previous results, CEBRA reliably distinguished risk assessment from escape behaviors using calcium activity in both *Vglut2*+ and *Vgat*+ populations across both tests (**Figure 4K-N, S3D-E**). Classification accuracy remained high across mice and was significantly above chance when compared to shuffled behavioral labels (**Figure 4L,N**), confirming that population activity robustly encodes behavioral state. Finally, we extended this approach by training CEBRA on all three threat conditions (**Fig S3F-G-H**). The model successfully distinguished both behavioral state and threat type with high accuracy, demonstrating that dPAG population activity contains sufficient information to decode not only the nature of the behavior, but also the context in which it occurs. As before, classification performance was cross-validated against shuffled labels, and accuracy dropped significantly in the control condition, confirming that the decoding was not driven by chance.

## Discussion

We have used *in vivo* single-unit calcium imaging to assess correlations between neural activity in glutamatergic and GABAergic neurons in dPAG and approach-escape behavior of laboratory mice towards predators, aggressive conspecifics and novel prey. Subsequently, we used optogenetic activation of GABAergic neurons in dPAG to investigate the causal relations between neural activity and approach-escape behavior. Two conclusions emerge from our findings. First, we found that, contrary to expectation, the two major firing cell classes previously described by *in vivo* single-unit electrophysiology studies in dPAG, *Assessment*+ and *Escape*+ neurons, exist amongst both glutamatergic and GABAergic neurons. This finding allows us to rule out the model in which the inverse activity patterns of *Assessment*+ and *Escape*+ neurons explained by their being, respectively, GABAergic and glutamatergic neurons. Instead, our data supports a model in which local GABAergic feedback inhibition between distinct sets of excitatory neurons promotes *Assessment*+ and *Escape*+ neurons. The feedback inhibition model is also compatible with the results of our optogenetic activation experiments. Second, we found a substantial overlap between *Assessment*+ and *Escape*+ neurons elicited by approach-avoidance to predator, aggressive conspecifics, and prey. This overlap argues for a convergence of approach-avoidance control encoding in dPAG rather than the persistence of parallel threat-class processing pathways, consistent with a role for this structure as a trigger for goal-directed defensive behavior regardless of the nature of the threat.

A notable paradox of our findings is that while GABAergic and glutamatergic dPAG neurons showed nearly indistinguishable neural firing correlates of approach and avoidance (both harbored *Assessment*+ and *Escape*+ cells), their optogenetic activation had opposite effects on defensive behavior. Optogenetic activation of glutamatergic neurons elicited freezing and flight (Tovote et al. 2016, Deng et al. 2016, Evans et al. 2018, Tsang et al. 2023), while optogenetic activation of GABAergic neurons inhibited defensive behaviors and promoted exploration (**Figure 3**; Tsang et al. 2023; Stempel et al. 2024). This discrepancy could be explained by a local circuit architecture in which glutamatergic neurons are projection neurons that promote defensive behaviors (Tsang et al., 2023), while GABAergic neurons are interneurons that mediate local feedback inhibition. In this model, GABAergic neurons would receive relatively sparse one-to-one excitatory inputs from local *Assessment*+ and *Escape*+ neurons and, as a result, themselves show *Assessment*+ or *Escape*+ firing activity. At the same time, optogenetic activation of glutamatergic neurons would promote defense via efferent pathways, while optogenetic activation of GABAergic interneurons would primarily inhibit principal glutamatergic neurons, suppressing defensive behaviors. A similar model has been proposed to explain *Assessment*+ and Flight+ neuron firing patterns in the medial hypothalamus (Rahy et al. 2022). Unfortunately, the local circuitry of dPAG remains largely unknown and current neural tracing and connectivity methods make it difficult to link *in vivo* neural activity with functional connectivity. However, local feedback inhibition could be tested by recording neural activity across glutamatergic and GABAergic neurons while optogenetically inhibiting one or the other population.

While the behavior correlations of glutamatergic and GABAergic neurons in dPAG were largely indistinguishable, some differences were observed. Although an assessment of their dynamics did not reveal statistically significant deviations during approach-avoidance behaviors (**Figure 2**), GABAergic neurons showed larger dynamics outside these behavior epochs than glutamatergic neurons and appeared to show tonic fluctuations in activity over longer timescales (**Figure 1, 2**), consistent with previous observations (Stempel et al., 2024). Unfortunately, we were not able to identify testing conditions under which the two cell types differed statistically, leaving open the question of whether GABAergic and glutamatergic neurons in dPAG reliably encode different aspects of defensive behavior. Our assessment of ensemble encoding of behavior also failed to shed light on cell-type specific differences in encoding, with both cell-types showing significant predictive correlations of approach and avoidance behaviors (**Figure 2, 4**).

Finally, our finding that the defensive behaviors to different threats are encoded in a common manner in dPAG is consistent with dPAG serving as a common pattern initiator (“trigger”) for stereotyped avoidance behaviors and previous manipulation studies showing that escape from predators and aggressive conspecifics both require an intact dPAG (Silva et al. 2013). Nevertheless, we note that the quality of approach-avoidance behaviors to each class of threat differed significantly. In particular, approach-avoidance bouts toward cockroaches were relatively short, and their duration was reflected in the corresponding neural activity dynamics, suggesting a direct relationship between dPAG activity dynamics and the quality or extent of defensive behavior (**Figure 4**). However, whether this relationship is causal remains unclear as we cannot rule out that there are efference copy feedback mechanisms by which dPAG activity is modulated by downstream motor systems. For example, we note that the quality of defensive behavior, including, for example, the direction of escape, is controlled downstream of dPAG (Tsang et al., 2023).

Our findings point to a local feedback circuitry in dPAG that is engaged by a wide variety of naturalistic threats and serves as a common trigger for defensive avoidance. The discovery of *Assessment*+ and *Escape*+ cells among neurons that both promote (glutamatergic) and inhibit (GABAergic) defense appears paradoxical and future work linking *in vivo* activity with circuit connectivity will be required to understand how this network produces innate freezing and flight behaviors. We believe that understanding this simple innate behavior initiation node will help understand the function of other brain structures where similar ON and OFF cell responses to behavior transitions have been observed.

## Materials and methods

### Animals

All experimental procedures involving the use of animals were carried out in accordance with the European Union (EU) Directive 2010/63/EU and under the approval of the EMBL Animal Use Committee and the Italian Ministry of Health License 541/2015-PR to Cornelius Gross. Adult mice (*Mus musculus*) were group housed in temperature and humidity-controlled cages with ad libitum access to food and water under a 12 h/12 h light–dark cycle. Mice in which optic fibers or headmounted GRIN lenses were implanted were singly housed after surgery. C57BL/6 J wild-type mice were obtained from local EMBL colonies. *Vglut2::Cre* and *Vgat::Cre* mice were obtained from JAX (stock no. 028863 and 028862), and were bred in-house on a C57BL/6J congenic background. The colonies were genotyped according to JAX recommendations for each transgenic line. Adult (>8 weeks old) transgenic mice in heterozygous state were used for experimental procedures. Only male mice were used in the experiments since they involved close social interactions with aggressive male mice. Aggressive mice used in the modified resident intruder test were male, ex-breeder adult CD1 mice obtained from local EMBL colonies or purchased from Charles Rivers Laboratories. CD1 mice were pre-screened for aggressiveness, and those attacking a C57BL/6J 3 or more times for three consecutive days were selected for further experiments. Adult male SHR/NHsd rats (*Rattus norvegicus*) purchased from Harlam were used as predators. Rats were singly housed in temperature and humidity controlled cages with ad libitum access to food and water under an inverted 12 h/12 h light–dark cycle. Turkestan cockroaches (*Blatta lateralis*) purchased from local commercial pet supply distributors were used as prey.

### Stereotaxic surgeries

Mice were anesthetized with 5% isoflurane and subsequently head-fixed in a stereotaxic frame (RWD Life Science) with body temperature maintained at 37 °C. Anesthesia was sustained with 1 to 2% isoflurane and oxygen. The skull was exposed, cleaned with hydrogen peroxide (0.3% in ddH2O) and leveled. Craniotomy was performed with a handheld drill. Either AAV5-hSyn::DIO-GCaMP6f (miniscope experiment), or AAV5-DIO-EF1a-hChR2(E123T/T159C)-EYFP (optogenetics experiment), or AAV5-DIO-EF1a-eYFP (optogenetics experiment control) was infused in the dPAG (4.30 posterior to bregma, 1.50 left to bregma and 2.65 mm ventral to the skull surface, at a 20° lateral angle). Injections were unilateral and ∼ 0.2 μl of virus was delivered with a pulled glass capillary (intraMARK, 10–20 μm tip diameter, Blaubrand). After 7-10’, the glass capillary was withdrawn and either a gradient-index optics lens (GRIN lens; Snap-imaging cannula model L type V, Doric Lenses) or an optic fiber attached to a ceramic ferrule (RWD, diameter 1.25 mm ceramic ferrule, 200 μm core, 0.22 NA, 3.3mm long) was implanted at a descending rate of 10μm/5-10s, at the same AP and ML coordinates but 100 μm above the virus injection site in the DV axis (2.55 mm ventral to the skull surface, at a 20° lateral angle). The implants were secured to the surface of the bone by a thin layer of first-layer cement (Superbond) followed by a layer of dental cement (Duralay). After the cement solidified, the skin was sutured and saline solution and Carprofen (5 mg/kg) were administered subcutaneously. Mice were allowed ca. 4 weeks for recovery and viral expression. pAAV.Syn.GCaMP6f.WPRE.SV40 was a gift from Douglas Kim & GENIE Project (Addgene viral prep # 100837-AAV5). pAAV-Ef1a-DIO hChR2(E123T/T159C)-EYFP and pAAV-Ef1a-DIO EYFP were gifts from Karl Deisseroth (Addgene viral prep # 35509-AAV5 and #27056-AAV5).

### Behavioral testing

#### Behavioral timeline

All behavioral testing was conducted in the mice’s light cycle. Miniscope recording experiments were carried out in male mice. The motivation for doing so was that social defeat is traditionally carried out in male-male dyads (Golden et al., 2011). To determine if PAG cells respond to threats in a specific or general manner, neurons need to be monitored longitudinally across exposure to different biological threats. To ensure this is possible with the miniscope system, mice were exposed, in a single session, to all threats. Mice were first exposed to the social threat, then to prey and last to the predator. Tests were distanced by a minimum of 5’ to allow the mouse and fluorescence to recover. Before behavioral testing, animals were habituated to handling, to the behavioral apparatus and to connection to wires for at least 5 days for the miniscope experiments and 3 days for optogenetics experiments. The behavioral arena was cleaned with water in between mice and with 70% ethanol at the end of the day.

#### Social exposure test

For mice to display defensive behaviors towards conspecifics, we first pre-selected aggressive CD1 male mice, and we had our experimental mice socially defeated twice. Aggressive CD1s were pre-selected if, in three consecutive days, they defeated different C57BL6/J male mice in a resident-intruder test, where CD1 mice were intruders in the C57BL6/J homecage. Social defeat was defined as 3-4 attacks in which CD1 chased and bit the C57BL6/J, which typically escaped, froze or displayed upright-postures.

**Social defeat:** social defeat sessions happened the two consecutive days immediately prior to the test day. (1) Assessment and escape phase: the experimental apparatus (custom made) consists of a chamber (“chamber”) attached to an elongated corridor (“corridor"). After enough time for the experimental animal to recover after connection to the miniscope wires, the CD1 was placed behind a wire mesh at the far end of the corridor respective to the chamber. The wire mesh permits interactions like sniffing and investigation but prevents defeat from happening. In this phase, the experimental mouse is free to assess and escape from the constrained CD1. (2) Defeat: the experimental mouse was constrained in the corridor and the CD1 released from the wire mesh for defeat to take place. Social defeat was stopped after 3-4 mild to moderate attacks. If a CD1 mouse did not attack, it was exchanged with another CD1. If the attacks were too aggressive (persistent biting and pulling the skin), defeat was stopped prematurely. (3) Postdefeat: after the CD1 mouse was removed from the apparatus, the experimental mouse stayed for another 5 minutes to consolidate the defeat experience. Importantly, mice encountered a different CD1 on each day of the test.

**Test day:** the test consisted of the assessment and escape phase described above. After two sessions of defeat, animals displayed repeated risk assessment and escape behavior and PAG activity was recorded with the miniscope during this phase of the test.

#### Prey exposure test

Mice were enclosed in the chamber and exposed to a freely moving cockroach. Mice displayed repeated risk assessment and escape behavior towards the cockroach. For more information regarding this behavioral test, see Rossier et al. 2021. In order to automatically estimate the distance to the cockroach, since they are free to move in the arena, the position of the cockroach was tracked by training a DeepLabCut (DLC) (Mathis et al., 2018) pose estimation pre-trained neural network.

#### Predator exposure test

A predator – a rat – is placed inside a second chamber connected with the far end of the corridor, with a glass separator with an aperture that permits sensory exchange but not aggressive contact. For more information regarding this test, see Masferrer et al. 2020.

#### Video capture and mouse tracking

All behavioral videos were recorded with a camera (Basler) providing a top view of the apparatus at either 25 or 40fps. X and y coordinates of 8 body parts the mouse were extracted by implementing a DLC (Mathis et al., 2018) pre-trained neural network. Afterward, for miniscope experiments, SimBA (Goodwin et al., 2024), a machine learning-based software for the classification of behavior, was used to correct outliers and to define the position of the animal in several regions of interest. A separate DLC model was trained to track the cockroach in exemplary videos. Custom-written Python scripts were used to calculate position-related metrics such as the velocity of the center of mass.

#### Manual behavioral annotation

Solomon Coder (http://solomoncoder.com/) was used for manual annotation of the following behaviors: (1) Risk assessment (slow, cautious approach towards the threat normally coupled with a stretch of the body) (2) Retraction (a very rapid retraction of the body, moving away from the aggressor), (3) Retreat (low-speed locomotion in the opposite direction of the threat, and towards a safe area – the chamber) and Flights (same as retreat, but high-speed locomotion). Unless specified, Escape behavior included retractions, retreats and flights.

### Miniscope experiments

#### System and synchronization with behavioral video

Animals in which GRIN lenses were implanted as described above were connected to a miniaturized fluorescent microscope or miniscope. The system (Doric Lenses) consists of a microscope body (Model L, Doric Lenses) that is attached to the GRIN lens implanted in the animal. This microscope receives light from a connectorized LED (458nm) through a 1m long mono optic fiber patch cord (MFP 200/230_900_0.48_0.8_FC_CM3, Doric Lenses). The miniscope is connected to a driver and ultimately to a computer through a 1.5mt long pigtailed electric cable (HDMI). An electric rotary joint (AHRJ OE_PT_ AH_12_HDMI, Doric Lenses) serves as a relay for both cables and prevents tangling. The system is operated with the DoricStudio software (Doric Lenses). Videos were recorded at 20 to 90% laser power and 100ms exposure time, which produced 10fps videos. Videos were stopped at a maximum length of 10 minutes to avoid photobleaching. Synchronization was ensured by the placement of an LED in the behavioral video FOV which was switched on by a TTL pulse sent by the miniscope driver when the miniscope was recording.

#### Miniscope video preprocessing

Videos were saved in .tif format by the DoricStudio software. They were pre-processed for background and motion correction following the below steps. Miniscope preprocessing and analysis scripts are available in our Gitlab repository (https://git.embl.org/grp-gross/excitatory_and_inhibitory_neurons_in_the_dorsal_periaqueductal_gray_encode_decisions_to_asses <u>s_and_escape_natural_threats</u>). (1) Scaling: a custom-written FiJi (Schindelin et al., 2012) macro was used to scale videos in the x and y dimensions from 630×630 to 315 by 315 pixels, significantly reducing the size of the videos which helped processing time, without compromising visual identification of neurons. (2) Scaled videos were screened for faulty frames, shown as full FOV back frames that occurred infrequently. Such frames were substituted by the mean of the previous and subsequent correct frame with a custom-written Python script. (3) Background subtraction: a FiJi macro was used to implement rolling-ball background subtraction algorithm. This algorithm performs a 3D spherical opening of the initial image, which can be visualized as follows: each video frame defines a 3D surface with height determined by its pixel intensity. A solid sphere moves constrained under this image surface. The background profile corresponds to the image surface that can be reached by the sphere, so high narrow peaks, whose diameter is smaller than the sphere’s diameter, are not included in the background’s profile. In this study we chose the sphere diameter to correspond to the typical size of a cell (Sternberg, 1983). (4) Motion correction: videos were motion corrected using the NoRMCorre algorithm (Pnevmatikakis & Giovannucci, 2017), which provides nonrigid registration of the video, that is particularly beneficial for miniscope recorded videos. NoRMCorre was implemented with CaImAn (Giovannucci et al., 2019) a toolbox including a variety of methods for the analysis of calcium imaging data. The provided Python scripts were adapted to allow for batch processing of our data in our HPC environment. Motion correction was run between 2 and 5 times to allow for optimal correction of shifts in XY. Algorithm parameters were optimized for each

| Round | 1 | 2 | 3 | 4 | 5 |
| --- | --- | --- | --- | --- | --- |
| gSig_filt | 8 |  |  |  |  |
| max_shifts | 20 | 10 |  |  |  |
| strides | 40 | 20 | 10 |  | 8 |
| overlaps | 20 | 10 | 5 |  | 4 |
| Max_deviation_rigid | 3 |  | 2 |  |  |

round of correction (table). Shifts in Z were not corrected.

#### ROI selection

Neurons were selected manually with the Fiji ROI manager by visual inspection of each video and its maximum projection. Overlapped neurons and those with large remaining shifts in the x and y axis were discarded. Mean fluorescence values over time were extracted for each ROI.

#### Normalization

Raw fluorescence traces were normalized by calculating *dF*/*F*0 and Z-score as follows:

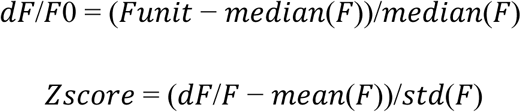

* Where F is a matrix of units x time (frames) and Funit is the activity of one neuron over time. For visualization purposes, data in heatmaps was normalized between -1 and 1.

#### Selection of responsive units

For each unit, we compared the average slope of its fluorescence trace in proximity to events of interest with a distribution of average slopes computed from random time points within the same time series.

More in details, given a fluorescence trace and a set of *N* time frames, the following steps were performed:

1. For each time frame *t*, take a window of length *L*=1s centered in *t*.
2. Align all windows and compute the average curve by averaging frame-by-frame
3. Compute the slope of the average curve as the coefficient of the line obtained by performing linear regression on the curve.
4. Repeat the following steps *n* times (e.g. *n* = 1000):

a. Extract *N* random frames from the time series.
b. For each random frame, consider a window of length *L* that centered on the frame.
c. Align all windows and compute the average time series with a frame-by-frame mean
d. Compute and store the slope of the curve as in point 3.
5. Use the stored "random" slopes to create a distribution by taking their histogram.
6. Compare the slope obtained at point 3. with the distribution to get its percentile value.

We defined a unit as responsive if its average fluorescence slope near behavioral events was “significant” compared to the random distribution. Specifically, a slope was considered significant if it was greater than the 0.975-quantile or smaller than the 0.025-quantile of the random distribution.

### Linear Time Warping

Linear time warping was employed to stretch or compress activity and behavioral trials along the time axis to match their median duration. Specifically, for each mouse and each unit, we first identified and stored sequences of behavioral events sorted in a specific order: “Leaving chamber”, “Assessment onset”, “Escape onset”, “Escape stop”. Sequences that included repetitions of one or more behaviors in consecutive spots were corrected by keeping only the last occurrence, e.g.[Leaving chamber, Assessment onset(1), Assessment onset(2), Escape onset, Escape stop] was corrected to [Leaving chamber, Assessment onset(2), Escape onset, Escape stop]. We then computed the median time gap between two consecutive behaviors over all stored trials (e.g. *m*_1_ = median time gap between Leaving chamber and Assessment onset) and used these values to subsequently perform linear interpolation on each trial. Specifically, consider for example the *j*-th activity trial of a neural activity curve *a*(*t*) with time frames *t*_1_, . . ., *t_k_*; the trial is then given by the neural activity values *a*(*t*_1_), . . ., *a*(*t_k_*). Consider *t*_!_ to be the time when behavior *A* occurs, *t_k_* the time when behavior *B* happens, and *m* to be the median duration of all trials of consecutive behaviors *A* and *B*. We divided the curve *a*(*t*) for trial *j* in *m* new equally spaced points *t*′_1_, . . ., *t*′*_m_* and estimated *a*(*t*′_1_), . . ., *a*(*t*′*_m_*) using linear interpolation. This procedure was applied to all trials of consecutive behaviors *A* and *B*, ensuring each had the same number of data points *m*.

### Population analysis with Cebra

Cebra (Schneider et al., 2023) is a computational framework based on nonlinear independent component analysis and contrastive learning to generate embeddings of high-dimensional neural data using auxiliary variables such as time or behavioral features. The model learns to distinguish between similar and dissimilar data points and it is penalized for producing similar embeddings for dissimilar points and vice versa. Cebra learns a mapping function from the high-dimensional training data to a lower dimensional space, in this study a three-dimensional sphere. Once trained, the same mapping function is applied to previously unseen test data to evaluate generalization. We employed multi-session Cebra Behavior using manually annotated behavioral labels as the auxiliary variable. To assess performance, we applied a K-nearest neighbors (KNN) classifier using cosine similarity (via scikit-learn). For each test data point in the low-dimensional space, we predicted its label based on the behavioral labels of its K=10 nearest training neighbors. The train-test split for multi-session Cebra Behavior was performed by randomly splitting each dataset without replacement using 70% of the data for the training set and by keeping the remaining 30% for testing. For each experimental test and for each mouse, a dataset was created by including the mouse’s neural activity as well as the manually annotated behavior, and by keeping only those data points where the mouse behavior was labeled as either ‘Flight’ or ‘Risk assessment’. We then used this data to build multiple models of multi-session Cebra Behavior; more specifically, we:

trained and tested Cebra on all datasets acquired during the predator experimental test. The same model was also tested on Vgat and Vglut mice data separately for the same experimental test. Two other CEBRA models were similarly trained and tested for the prey and social datasets.trained and tested a CEBRA models on the Vgat mice data recorded during the predator experimental test. We repeated a similar analysis only on the Vglut mice for the same experimental test.concatenated the three datasets acquired during the different experimental tests for each single mouse after normalizing them (z-score). This time, the behavioral features were transformed to include the information about the experimental test (e.g. ‘Flight’ was transformed into ‘Flight_prey’ for the prey test and ‘Flight_social’ for the social test), resulting in six-class categorical features. We then trained and tested Cebra on the resulting datasets. The same model was also tested on *Vglut2* and *Vgat* mice data separately. All experiments above were repeated after randomly shuffling behavioral labels.

In all cases, we trained Cebra Behavior with the default parameters except for: model_architecture = “offset10-model”, batch_size = 512, learning_rate = 0.001, output_dimension = 3.

### Optogenetic experiments

#### Optogenetic stimulation parameters

Optogenetic experiments were carried in male (n=4) and female (n=4) male mice in the rat exposure test. Blue (465 nm) laser light (PSU-III-LED, CNI laser) was delivered in alternated minutes for a total of 6 minutes. Laser light reached the optic fiber ferrules through a rotary joint (Thorlabs) and high-performance bilateral pathcords (Thorlabs). Light intensity at the tip of the patch cord was measured with a lightmeter (Thorlabs) before and after each test session and set to 15-20mWatt. Light was delivered in 10ms pulses and stimulation trains were programmed and generated with Pulser Software. eYFP and control animals were tested with the same light stimulation protocol. Experimenters were blind to genotype during experimentation and analysis.

#### Video acquisition and tracking

Behavioral videos were recorded with a top camera (Basler) at 25 fps. X and y coordinates of 16 body parts of the mouse were tracked with a custom-trained DeepLabCut (Mathis et al., 2018) model and outlier corrected with SimBA (Goodwin et al., 2024). Custom-written Python scripts were used to calculate position-related metrics such as the velocity of the center of mass.

#### Unsupervised behavioral annotation

Keypoint-MoSeq (Weinreb et al., 2024) was implemented for the unbiased categorization of behavior in syllables based on 16 bodypart DeepLabCut (Mathis et al., 2018) extracted x and y coordinates. Selected syllables were matched with manually annotated behaviors by inspecting the software-provided videos and by the analysis of the overlap of automatically and manually annotated behavior. Behavior definitions: (1) Risk assessment: stretch attend or stretch approach postures, directed towards threat source. (2) Retraction: rapid retraction of the head. (3) Escape: high-velocity locomotion aiming at increasing distance between threat source and subject. (4) Rearing: standing on hindpaws with forepaws leaning to the walls of the behavioral box. (5) Jumping: rapid upright motion towards top of the behavioral box.

## Acknowledgements

We thank the EMBL Rome Laboratory Animal Facility, Gene Editing & Virus Facility, Microscopy Facility, and Roberto Voci and Valerio Rossi for support with animal husbandry and management. We thank Maria E. Masferrer for support with data collection and analysis. The work was funded by the European Molecular Biology Laboratory (EMBL).

## Author contributions

I.P.A. and C.T.G. designed research; I.P.A., V.M. and T.S. performed research; I.P.A and S.T. analyzed data; and I.P.A., S.T. and C.T.G. wrote the paper.

## References

Amano, K., Tanikawa, T., Kawamura, H., Iseki, H., Notani, M., Kawabatake, H., Shiwaku, T., Suda, T., Demura, H., & Kitamura, K. (1982). Endorphins and pain relief. Further observations on electrical stimulation of the lateral part of the periaqueductal gray matter during rostral mesencephalic reticulotomy for pain relief. Applied neurophysiology, 45(1-2), 123–135.

Beitz, A. J., (1982). The organization of afferent projections to the midbrain periaqueductal gray of the rat. Neuroscience, 7(1), 133–159. 10.1016/0306-4522(82)90157-9

Bittencourt, A. S., Carobrez, A. P., Zamprogno, L. P., Tufik, S., & Schenberg, L. C. (2004). Organization of single components of defensive behaviors within distinct columns of periaqueductal gray matter of the rat: role of N-methyl-D-aspartic acid glutamate receptors. Neuroscience, 125(1), 71–89. 10.1016/j.neuroscience.2004.01.026

Canteras, N. S. (2002). The medial hypothalamic defensive system: Hodological organization and functional implications. Pharmacology Biochemistry and Behavior, 71(3), 481–491. 10.1016/S0091-3057(01)00685-2

Carrive P. (1993). The periaqueductal gray and defensive behavior: functional representation and neuronal organization. Behavioural brain research, 58(1-2), 27–47. 10.1016/0166-4328(93)90088-8

Deng, H., Xiao, X., & Wang, Z. (2016). Periaqueductal Gray Neuronal Activities Underlie Different Aspects of Defensive Behaviors. The Journal of Neuroscience, 36(29), 7580–7588. 10.1523/JNEUROSCI.4425-15.2016

Esteban Masferrer, M., Silva, B., Nomoto, K., Lima, S., & Gross, C. (2020). Differential Encoding of Predator Fear in the Ventromedial Hypothalamus and Periaqueductal Grey. TThe Journal of Neuroscience, 40(48), 9283–9292. 10.1523/JNEUROSCI.0761-18.2020

Evans, D. A., Stempel, A. V., Vale, R., Ruehle, S., Lefler, Y., & Branco, T. (2018). A synaptic threshold mechanism for computing escape decisions. Nature, 558(7711), 590–594. 10.1038/s41586-018-0244-6

Fillinger, C., Yalcin, I., Barrot, M., & Veinante, P. (2018). Efferents of anterior cingulate areas 24a and 24b and midcingulate areas 24a’ and 24b’ in the mouse. Brain structure & function, 223(4), 1747–1778. 10.1007/s00429-017-1585-x

Giovannucci, A., Friedrich, J., Gunn, P., Kalfon, J., Brown, B. L., Koay, S. A., Taxidis, J., Najafi, F., Gauthier, J. L., Zhou, P., Khakh, B. S., Tank, D. W., Chklovskii, D. B., & Pnevmatikakis, E. A. (2019). CaImAn an open source tool for scalable calcium imaging data analysis. ELife, 8. 10.7554/eLife.38173

Goodwin, N. L., Choong, J. J., Hwang, S., Pitts, K., Bloom, L., Islam, A., Zhang, Y. Y., Szelenyi, E. R., Tong, X., Newman, E. L., Miczek, K., Wright, H. R., McLaughlin, R. J., Norville, Z. C., Eshel, N., Heshmati, M., Nilsson, S. R. O., & Golden, S. A. (2024). Simple Behavioral Analysis (SimBA) as a platform for explainable machine learning in behavioral neuroscience. Nature Neuroscience 2024 27:7, 27(7), 1411–1424. 10.1038/s41593-024-01649-9

Gross, C. T., & Canteras, N. S. (2012). The many paths to fear. Nature Reviews Neuroscience, 13(9), 651–658. 10.1038/nrn3301

Helassa, N., Podor, B., Fine, A., & Török, K. (2016). Design and mechanistic insight into ultrafast calcium indicators for monitoring intracellular calcium dynamics. Scientific Reports 2016 6:1, 6(1), 1–14. 10.1038/srep38276

Johansen, J. P., Tarpley, J. W., LeDoux, J. E., & Blair, H. T. (2010). Neural substrates for expectation-modulated fear learning in the amygdala and periaqueductal gray. Nature neuroscience, 13(8), 979–986. 10.1038/nn.2594

Marchand, J. E. and Hagino, N. (1983), Afferents to the periaqueductal gray in the rat. A horseradish peroxidase study. Neuroscience, 9(1):95–106. 10.1016/0306-4522(83)90049-0.

Mathis, A., Mamidanna, P., Cury, K. M., Abe, T., Murthy, V. N., Mathis, M. W., & Bethge, M. (2018). DeepLabCut: markerless pose estimation of user-defined body parts with deep learning. Nature Neuroscience, 21(9), 1281–1289. 10.1038/s41593-018-0209-y

Motta, S. C., Gouveia, F. V., Baldo, M. V. C., Canteras, N. S., & Swanson, L. W. (2009). Dissecting the brain’s fear system reveals the hypothalamus is critical for responding in subordinate conspecific intruders. Proceedings of the National Academy of Sciences of the United States of America, 106(12), 4870–4875. 10.1073/pnas.0900939106

Nashold, B. S., Wilson, W. P., & Slaughter, D. G. (1969). Sensations evoked by stimulation in the midbrain of man. Journal of Neurosurgery, 30(1), 14–24. 10.3171/jns.1969.30.1.0014

Pnevmatikakis, E. A., & Giovannucci, A. (2017). NoRMCorre: An online algorithm for piecewise rigid motion correction of calcium imaging data. Journal of Neuroscience Methods, 291, 83–94. 10.1016/J.JNEUMETH.2017.07.031

Rahy, R., Asari, H., & Gross, C. T. (2022). Sensory-thresholded switch of neural firing states in a computational model of the ventromedial hypothalamus. Frontiers in Computational Neuroscience, 16, Article 813132. 10.3389/fncom.2022.813132

Reis, F. M. C. V., Liu, J., Schuette, P. J., Lee, J. Y., Maesta-Pereira, S., Chakerian, M., Wang, W., Canteras, N. S., Kao, J. C., & Adhikari, A. (2021). Shared dorsal periaqueductal gray activation patterns during exposure to innate and conditioned threats. The Journal of Neuroscience, 41(25), 5399–5420. 10.1523/JNEUROSCI.2450-20.2021

Reis, F. M. C. V., Lee, J. Y., Maesta-Pereira, S., Schuette, P. J., Chakerian, M., Liu, J., La-Vu, M. Q., Tobias, B. C., Ikebara, J. M., Kihara, A. H., Canteras, N. S., Kao, J. C., & Adhikari, A. (2021). Dorsal periaqueductal gray ensembles represent approach and avoidance states. eLife, 10, Article e64934. 10.7554/eLife.64934

Rossier, D., La Franca, V., Salemi, T., Natale, S., Gross, C. T., Franca, V. La, Salemi, T., Natale, S., & Gross, C. T. (2021). A neural circuit for competing approach and defense underlying prey capture. Proceedings of the National Academy of Sciences, 118(15), e2013411118. https://www.pnas.org/content/118/15/e2013411118

Schindelin, J., Arganda-Carreras, I., Frise, E., Kaynig, V., Longair, M., Pietzsch, T., Preibisch, S., Rueden, C., Saalfeld, S., Schmid, B., Tinevez, J.-Y., White, D. J., Hartenstein, V., Eliceiri, K., Tomancak, P., & Cardona, A. (2012). Fiji: an open-source platform for biological-image analysis. Nature Methods 2012 9:7, 9(7), 676–682. 10.1038/nmeth.2019

Schneider, S., Lee, J. H., & Mathis, M. W. (2023). Learnable latent embeddings for joint behavioural and neural analysis. Nature, 617(7960), 360–368. 10.1038/s41586-023-06031-6

Silva, B. A., Mattucci, C., Krzywkowski, P., Murana, E., Illarionova, A., Grinevich, V., Canteras, N. S., Ragozzino, D., & Gross, C. T. (2013). Independent hypothalamic circuits for social and predator fear. Nature Neuroscience, 16(12), 1731–1733. 10.1038/nn.3573

Stempel, A. V., Evans, D. A., Arocas, O. P., Claudi, F., Lenzi, S. C., Kutsarova, E., Margrie, T. W., & Branco, T. (2024). Tonically active GABAergic neurons in the dorsal periaqueductal gray control instinctive escape in mice. Current Biology, 34(13), 3031–3039.e7. 10.1016/J.CUB.2024.05.068/ATTACHMENT/C5BB0710-CE30-4201-A55E-D0DB0A110409/MMC6.PDF

Sternberg, S. R. (1983). Biomedical Image Processing. Computer, 16(1), 22–34. 10.1109/MC.1983.1654163

Tovote, P., Esposito, M. S., Botta, P., Chaudun, F., Fadok, J. P., Markovic, M., Wolff, S. B. E., Ramakrishnan, C., Fenno, L., Deisseroth, K., Herry, C., Arber, S., & Lüthi, A. (2016). Midbrain circuits for defensive behaviour. Nature, 534(7606), 206–212. 10.1038/nature17996

Tsang, E., Orlandini, C., Sureka, R., Crevenna, A. H., Perlas, E., Prankerd, I., Masferrer, M. E., & Gross, C. T. (2023). Induction of flight via midbrain projections to the cuneiform nucleus. PloS One, 18(2), e0281464. 10.1371/JOURNAL.PONE.0281464

Vaughn, E., Eichhorn, S., Jung, W., Zhuang, X., & Dulac, C. (2022). Three-dimensional Interrogation of Cell Types and Instinctive Behavior in the Periaqueductal Gray. BioRxiv, 2022.06.27.497769. 10.1101/2022.06.27.497769

Vianna, D. M., & Brandão, M. L. (2003). Anatomical connections of the periaqueductal gray: specific neural substrates for different kinds of fear. Brazilian journal of medical and biological research = Revista brasileira de pesquisas medicas e biologica, 36(5), 557–566. 10.1590/s0100-879x2003000500002

Weinreb, C., Pearl, J. E., Lin, S., Abdal, M., Osman, M., Zhang, L., Annapragada, S., Conlin, E., Hoffmann, R., Makowska, S., Gillis, W. F., Jay, M., Ye, S., Mathis, A., Mathis, M. W., Pereira, T., Linderman, S. W., Sandeep, &, & Datta, R. (2024). Keypoint-MoSeq: parsing behavior by linking point tracking to pose dynamics. Nature Methods 2024 21:7, 21(7), 1329–1339. 10.1038/s41592-024-02318-2

Wu, G.-Y., Li, R.-X., Liu, J., Sun, L., Yi, Y.-L., Yao, J., Tang, B.-Q., Wen, H.-Z., Chen, P.-H., Lou, Y.-X., Li, H.-L., & Sui, J.-F. (2024). An excitatory neural circuit for descending inhibition of itch processing. Cell Reports, 43(12), Article 115062. 10.1016/j.celrep.2024.115062

