## Supplementary materials for "Excitatory and inhibitory neurons in the dorsal periaqueductal gray encode decisions to assess and escape natural threats"

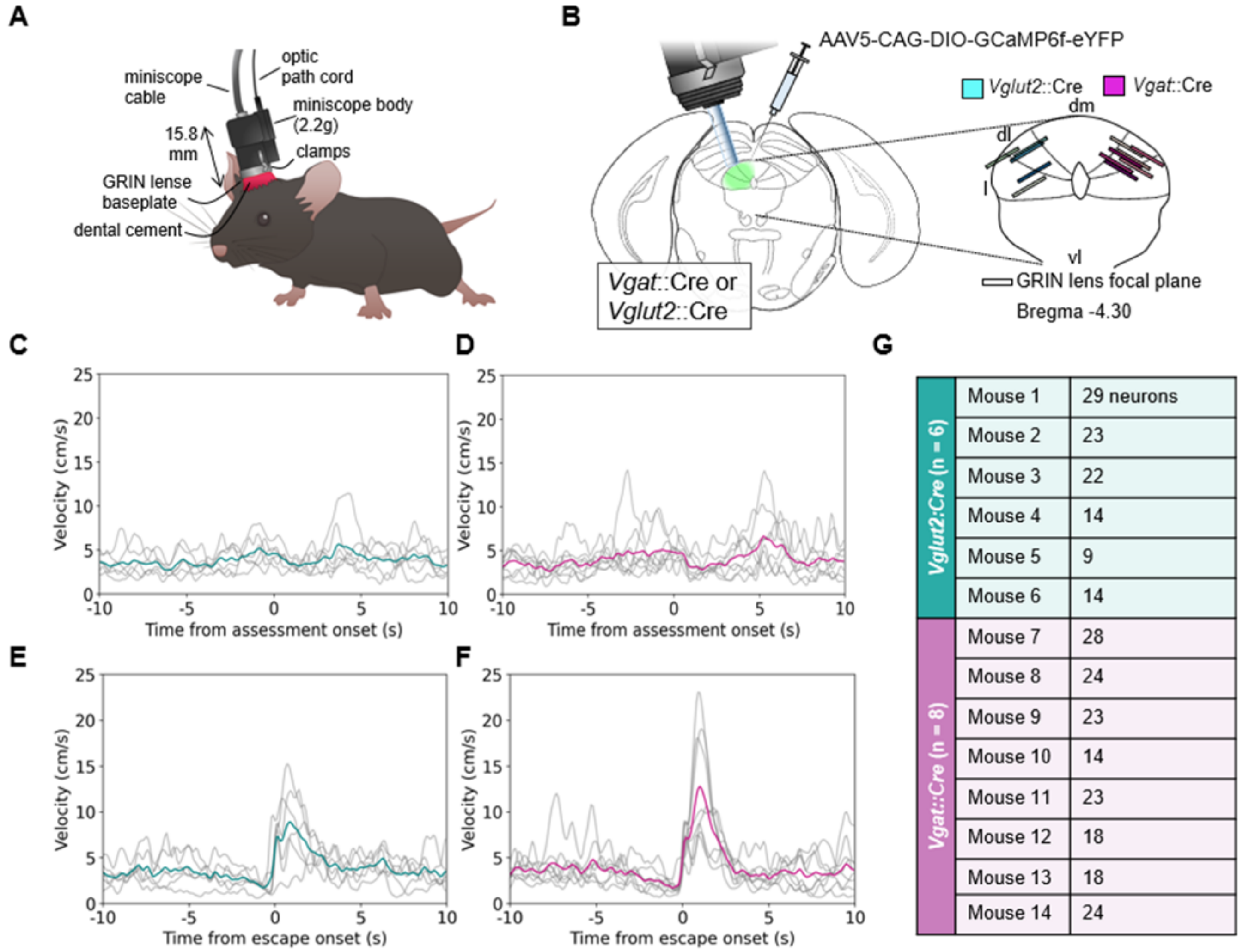


***Supplementary Figure 1*. Further information on GRIN lens placements, behavior and FOVs.** (**A**) Schematic of the miniscope system. (**B**) Summary of GRIN lens placement for all *Vglut2*::Cre and *Vgat*::Cre mice. (**C**) Velocity of the center of mass centered around assessment onset for *Vglut2*::Cre mice. (**D**) Velocity of the center of mass centered around assessment onset for *Vgat*::Cre mice. (**E**) Velocity of the center of mass centered around escape onset for *Vglut2*::Cre mice. (**F**) Velocity of the center of mass centered around escape onset for *Vgat*::Cre mice. (**G**) Number of observed neurons per field of view.

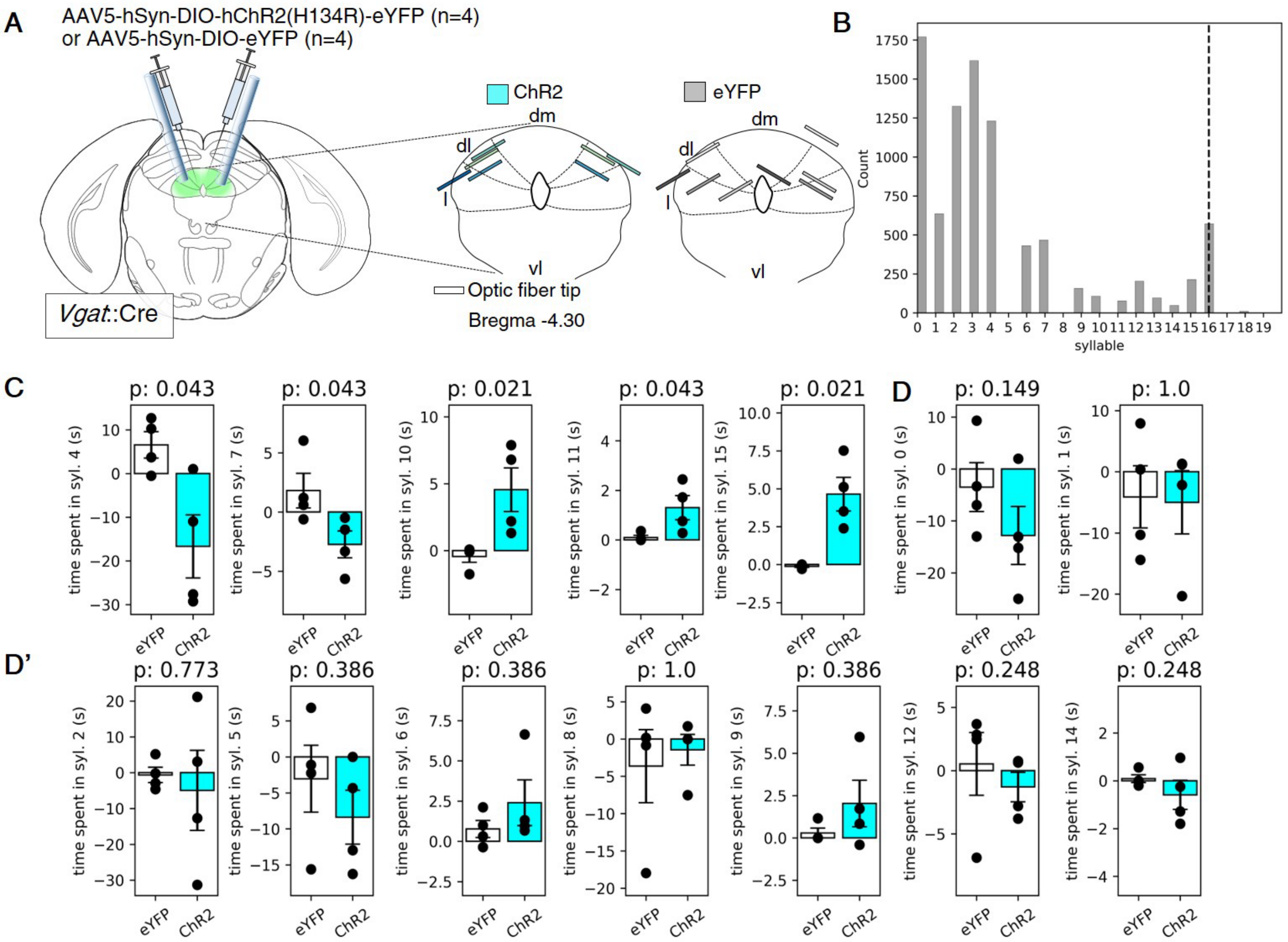


***Supplementary Figure 2*. Optogenetic stimulation of *Vgat*+ dPAG neurons modulates defensive and exploratory behavior.** (**A**) Summary of optic fiber placement for eYPF (grey) and ChR2 (cyan) mice. (**B**) Distribution of frequencies of kp-moSeq detected syllables. The dashed line indicates 95% of the data. (**C**) Light-induced difference in time spent in kp-moSeq syllables for eYFP and ChR2 mice that tested significant by the Wilcoxon rank-sum test, or (**D** and **D’**) non-significant.

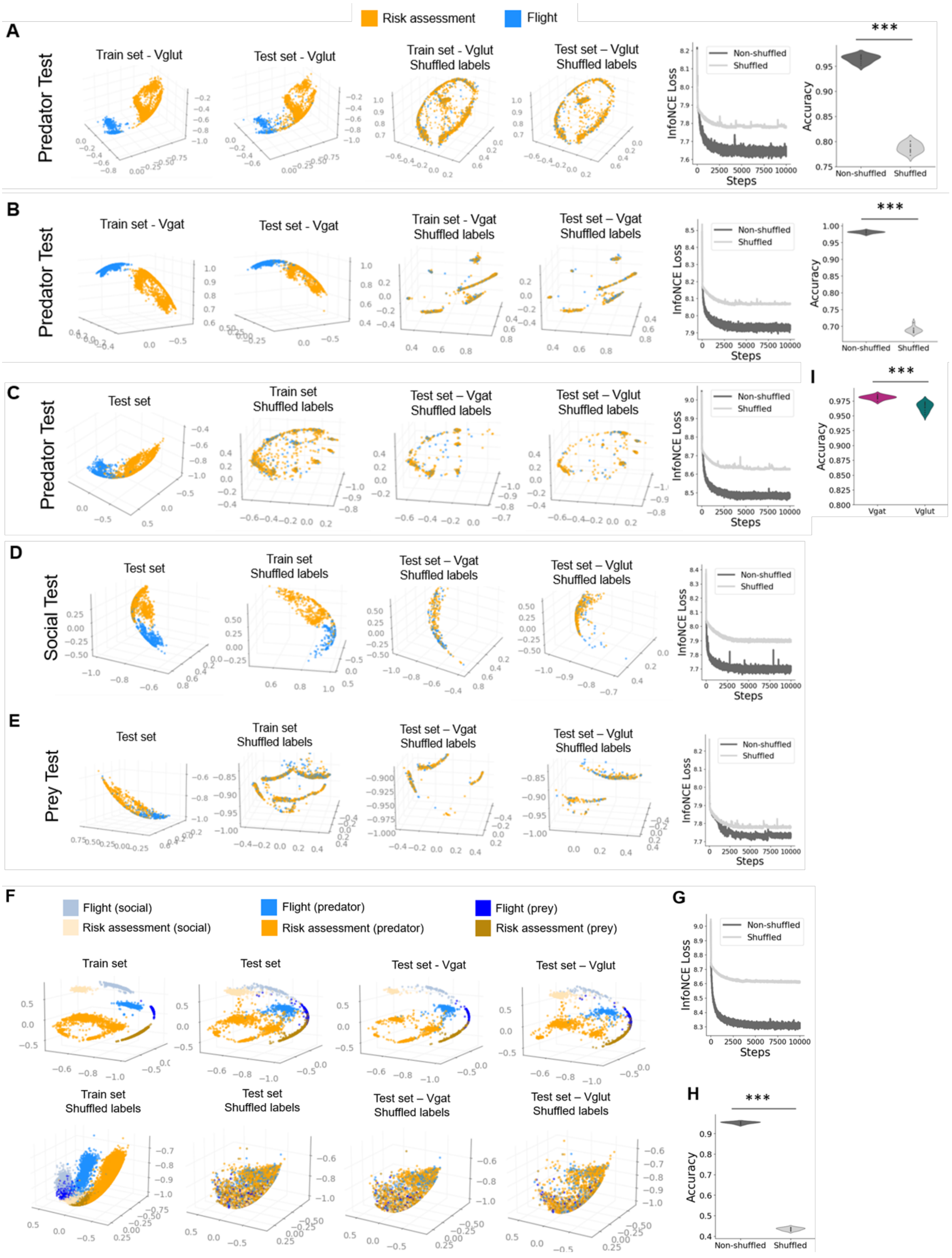


***Supplementary Figure 3.*** (**A**) From left to right: Train embedding obtained by training multi-session Cebra Behavior on 70% of each Vglut mouse's neural activity and behavioral labels acquired during the predator experimental test; test embedding obtained by testing the previous model on unseen data acquired from Vglut mice during the same experimental test; multi-session Cebra Behavior train embedding obtained by training the model on 70% of each Vglut mouse's neural activity and behavioral labels acquired during the predator experimental test after randomly shuffling the behavioral labels; test embedding obtained by testing the previous model on unseen data with shuffled behavioral labels acquired from Vglut mice during the same experimental test; comparison of the InfoNCE loss functions for the two non-shuffled and shuffled multi-session Cebra Behavior models trained and tested using only Vglut data from the predator test; comparison between prediction accuracy of behavioral labels using KNN of 1) 10 different multi-session Cebra Behavior models trained and tested on 10 separate random train-test splits (Vglut mice, predator test) and 2) 10 different multi-session Cebra Behavior models trained and tested with shuffled behavioral labels (same mice, predator test, new random train-test split). Statistical significance was tested with Wilcoxon rank-sum, ***$p < 0.001$, **$p < 0.01$, *$p < 0.05$. (**B**) As in (A) but for Vgat mice only. (**C**) From left to right: Test embedding obtained from multi-session Cebra Behavior trained on 70% of each mouse's neural activity and behavioral labels acquired during the predator experimental test; train embedding obtained from multi-session Cebra Behavior trained on 70% of each mouse's neural activity and behavioral labels acquired during the predator experimental test after randomly shuffling the behavioral labels; test embedding obtained by testing the previous model on unseen data acquired from Vgat mice during the same experimental test; test embedding similarly obtained by testing the previous model on Vglut data from the same experimental test; test embedding obtained by testing the previous model on all data (Vglut + Vgat mice) acquired during the same experimental test; comparison of the InfoNCE loss functions for the two non-shuffled and shuffled multi-session Cebra Behavior models trained and tested on all data from the predator test. (**D**) As in (C) but for the social experimental test (**E**) As in (C) but for the prey experimental test (**F**) First row, from left to right: train embedding obtained by training multi-session Cebra Behavior on 70% of each mouse' normalized neural activity and behavioral labels acquired during all experimental tests; test embedding obtained by testing the previous model on unseen data acquired from all mice during all experimental tests; test embedding obtained by testing the previous model on unseen data acquired from Vgat mice during all experimental tests; test embedding similarly obtained by testing the previous model on Vglut data from all experimental tests. Second row, from left to right: train embedding obtained by training multi-session Cebra Behavior on 70% of each mouse' normalized neural activity and behavioral labels acquired during all experimental tests after randomly shuffling the behavioral labels; test embedding obtained by testing the previous model on unseen data acquired from all mice during all experimental tests; test embedding obtained by testing the previous model on unseen data acquired from Vgat mice during all experimental tests; test embedding similarly obtained by testing the previous model on Vglut data from all experimental tests; (**G**) comparison of the InfoNCE loss functions for the two non-shuffled and shuffled multi-session Cebra Behavior models trained and tested using all data from all experimental tests as in (F). (**H**) Comparison between prediction accuracy of behavioral labels using KNN of 1) 10 different multi-session Cebra Behavior models trained and tested on 10 separate random train-test splits (all mice, all tests) and 2) 10 different multi-session Cebra Behavior models trained and tested with shuffled behavioral labels (same mice, all test, new random train-test split). (**I**) Prediction accuracy of behavioral labels using KNN of 10 different multi-session Cebra Behavior models trained and tested on 10 different randomly extracted train and test data from Vgat and Vglut mice separately. Statistical significance was tested with Wilcoxon rank-sum, ***$p < 0.001$, **$p < 0.01$, *$p < 0.05$
